# Shedding light on the grey zone of speciation along a continuum of genomic divergence

**DOI:** 10.1101/059790

**Authors:** Camille Roux, Christelle Fraïsse, Jonathan Romiguier, Yoann Anciaux, Nicolas Galtier, Nicolas Bierne

## Abstract

Speciation results from the progressive accumulation of mutations that decrease the probability of mating between parental populations, or reduce the fitness of hybrids - the so-called species barriers. The speciation genomic literature, however, is mainly a collection of case studies, each with its own approach and specificities, such that a global view of the gradual process of evolution from one to two species is currently lacking. Of primary importance is the prevalence of gene flow between diverging entities, which is central in most species concepts, and has been widely discussed in recent years. Here we explore the continuum of speciation thanks to a comparative analysis of genomic data from 61 pairs of populations/species of animals with variable levels of divergence. Gene flow between diverging gene pools is assessed under an Approximate Bayesian Computation (ABC) framework. We show that the intermediate "grey zone" of speciation, in which taxonomy is often controversial, spans from 0.5% to 2% of net synonymous divergence, irrespective of species life-history traits or ecology. Thanks to appropriate modeling of among-loci variation in genetic drift and introgression rate, we clarify the status of the majority of ambiguous cases and uncover a number of cryptic species. Our analysis also reveals the high incidence in animals of semi-isolated species, when some but not all loci are affected by barriers to gene flow, and highlights the intrinsic difficulty, both statistical and conceptual, of delineating species in the grey zone of speciation.

## Introduction

An important issue in evolutionary biology is understanding how the continuous-time process of speciation can lead to discrete entities - species. There is usually no ambiguity about species delineation when distant lineages are compared. The continuous nature of the divergence process, however, causes endless debates about the species status of closely-related lineages [1]. A number of definitions of species have thus been introduced over the 20th century, each of them using its own criteria - morphological, ecological, phylogenetic, biological, evolutionary or genotypic. A major problem is that distinct markers do not diverge in time at the same rate [2]. For instance, in some taxa, morphological differences evolve faster than the expression of hybrid fitness depression, which in turn typically establishes long before genome-wide reciprocal monophyly [3]. In other groups, morphology is almost unchanged between lineages that show high levels of molecular divergence [4]. The erratic behaviour and evolution of the various criteria is such that in a wide range of between-lineage divergence, named the grey zone of the speciation continuum, distinct species concepts do not converge to the same conclusions regarding species delineation [2].

Besides taxonomic aspects, the grey zone has raised an intense controversy regarding the genetic mechanisms involved in the formation of species [5–7]. Of peculiar importance is the question of gene flow between diverging lineages. How isolated two gene pools must be for speciation to begin? How long does gene flow persist as lineages diverge? Is speciation a gradual process of gene flow interruption, or a succession of periods of isolation and periods of contact? These questions are not only central in the speciation literature, but also relevant to the debate about species delineation, the ability of individuals to exchange genes being at the heart of the biological concept of species.

As genomic data have become easier and less expensive to obtain, sophisticated computational approaches have been developed to perform historical inferences in speciation genomics, *i.e.*, estimate the time of ancestral separation in two gene pools, changes in effective population size over evolutionary times, and the history of gene flow between the considered lineages [8–10]. Simulation-based Approximate Bayesian Computation (ABC) methods are particularly flexible and have recently attracted an increased attention in speciation genomics. One strength of ABC approaches is their ability to deal with complex, hopefully realistic models of speciation, and test for the presence or absence of ongoing introgression between sister lineages. This is achieved by simulating molecular data under alternative models of speciation with or without current introgression, and choosing among models based on their relative posterior probabilities [11].

Migration tends to homogenize allele content and frequency between diverging populations. This homogenizing effect, however, is often expected to only affect a fraction of the genome. This is because the effective migration rate is impeded in regions containing loci involved in assortative mating, hybrid fitness depression, or other mechanisms of isolation — the so-called genetic barriers [12]. Consequently, gene flow is best identified by models explicitly accounting for among-loci heterogeneity in introgression rates, as demonstrated by a number of recent studies [13–16]. When homogeneous introgression rate across the genome is assumed, distant lineages having accumulated a large number of genetic barriers can be inferred as currently isolated whereas they actually exchange alleles at a minority of loci unlinked to barriers [14]. Conversely, closely related lineages can be inferred as currently exchanging genes while some regions of the genome are already evolving independently [16,17] such that heterogeneous introgression models can provide support to the genic view of speciation [18]. Besides, introgression rates alone do not govern local patterns of genetic differentiation [19]. Directional selective processes, such as hitch-hiking effects [20] or background selection [21], are expected to affect the landscape of population differentiation by lowering polymorphism levels at particular loci, especially in low recombining or gene-dense genomic regions. Neglecting this confounding effect tends to inflate the proportion of false-positives in statistical tests of ongoing gene flow [19] and to mislead inferences [22]. Linked directional selection is expected to locally increase the stochasticity of allele frequency evolution, a process sometimes coined genetic draft [23]. Its effect can therefore be modeled by assuming that the effective population size, *Ne*, which determines the strength of genetic drift, varies among loci [24].

Multi-locus analyses of the process of population divergence has been achieved in various groups of animals [25,26] and plants [27–29] in which genome-wide data are available, revealing a diversity of patterns. These case studies, however, are limited in number and have taken different approaches, so that we still lack a unifying picture of the prevalence of gene flow during early divergence between gene pools. Here, we gathered a dataset of 61 pairs of populations/species of animals occupying a wide continuum of divergence level. Species were selected in order to sample the phylogenetic and ecological diversity of animals [30], irrespective of any aspect related to population structure or speciation. We investigated the effects of genomic divergence between populations on patterns of gene flow, paying attention to the ability of ABC methods to distinguish between competing models and the influence of model assumptions.

## Results

### Simulations: ABC as a powerful approach to test for current introgression

Five demographic models differing by the assumed history of gene flow were considered (Fig. 1), namely strict isolation (SI), ancient migration (AM), isolation with migration (IM), secondary contact (SC) and panmixia (PAN). The later three models involve ongoing gene flow between the two studied populations, whereas the former two do not. The five demographic models were subdivided into different genomic submodels, which reflect alternative assumptions about the genomic distribution of indirect selective effects on the effective population size (HomoN if homogeneous or HeteroN if heterogeneous) and on the migration rate (HomoM if homogeneous or HeteroM if heterogeneous). Heterogeneous effective population size was considered in all the models while heterogeneous migration rate was considered in models with gene flow (IM, AM and SC). The SI and PAN models were divided into two submodels (HomoN and HeteroN) and the AM, IM and SC models were divided into four submodels (HomoN_HomoM, HomoN_HeteroM, HeteroN_HomoM and HeteroN_HeteroM).

**Figure 1.**
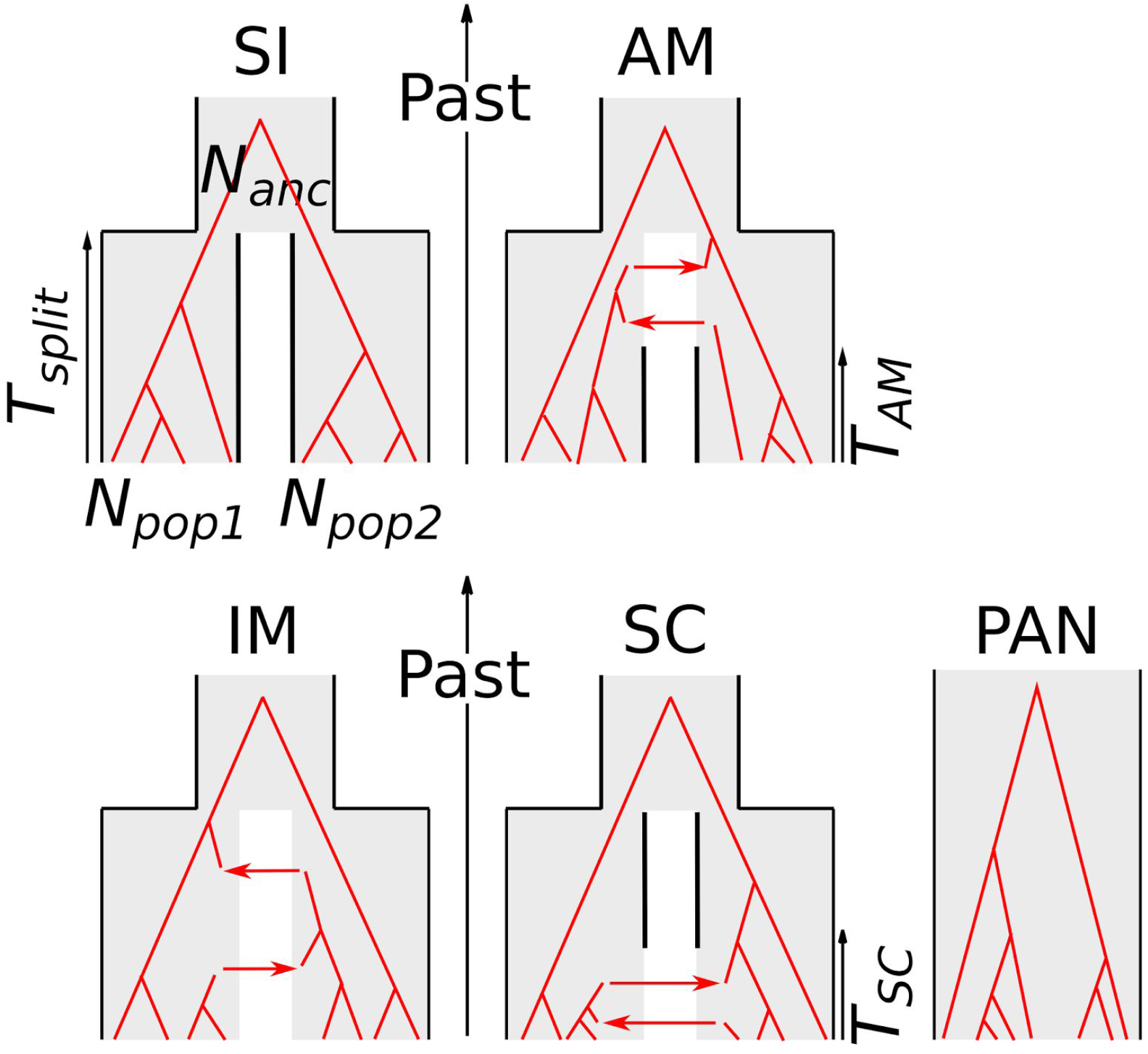
Compared alternative models of speciation. SI=Strict Isolation: subdivision of an ancestral diploid panmictic population (of size *N*_*anc*_) in two diploid populations (of constant sizes *N*_*pop1*_ and *N*_*pop2*_) at time *T*_*split\**_. AM=Ancestral Migration: the two newly formed populations continue to exchange alleles until time *T*_*AM*_. IM= Isolation with Migration: the two daughter populations continuously exchange alleles until present time. SC=Secondary Contact: the daughter populations first evolve in isolation then experience a secondary contact and start exchanging alleles at time *T*_SC_. PAN: Panmictic model. All individuals are sampled from the same panmictic population. Red phylogenies represent possible gene trees under each alternative models.

The dominant assumption in published demographic inferences is the HomoN submodel, in which it is assumed that most of the genetic variation in the genome is unaffected, or equally affected, by selection at linked sites. Here, HomoN was simulated using a single value of effective population size shared by all loci across genome, but the effective population size differed among populations. The HeteroN submodel accounts for local genomic effects of directional selection (background selection, selective sweeps) by considering a variable effective population size among loci, here assumed to follow a rescaled Beta distribution. The HomoM submodel assumes that all loci share the same probability to receive alleles from the sister population, i.e., posits the absence of species barrier or of adaptively introgressed locus. Alternatively, the HeteroM submodel accounts for the existence of local barriers to gene flow, of variable strengths, and of variable levels of genetic linkage to the sampled loci. HeteroM was here simulated by assuming that the effective introgression rate is Beta distributed across the genome, thus intending to account for the combined effects of selection, recombination and gene density. In principle one could explicitly include information on local recombination rates and gene density but no such data was available in the species of our dataset.

We explicitly tested the hypothesis of current gene flow by comparing the relative posterior probabilities of 16 models for 61 pairs of species distributed along a continuum of molecular divergence. In the ABC framework, the posterior probability of a model corresponds to its relative ability to theoretically produce datasets similar to the observed dataset, compared to a set of alternative models. Before analyzing datasets from the 61 pairs of animal species, we first assessed the power of the adopted ABC approach to correctly distinguish between models involving current isolation (SI + AM) *vs*. ongoing migration (IM + SC + PAN). This was achieved by randomly simulating 116,000 datasets distributed over the 16 compared models and applying our ABC inference method to each of them. Specifically, we investigated which model had the highest posterior probability and assessed significance by estimating the associated robustness, the probability to correctly support a model given its posterior probability. A robustness greater than 0.95 can be interpreted as a *P* value below 0.05 [31]. The analysis of simulated datasets allowed us to empirically measure a threshold value of 0.6419 for the posterior probability *P*_migration_ (= *P*_IM_+*P*_SC_+*P*_PAN_), above which the robustness to support ongoing migration is greater than 0.95. Similarly, a posterior probability *P*_migration_ below 0.1304 implied a statistical support for the current isolation model with a robustness greater than 0.95.

Among the 58,000 simulated datasets in which current gene flow was assumed (IM, SC, PAN; Fig. 2-A), 99.462% were true positives (*P*_migration_ > *P*_isolation_ and robustness ≥ 0.95), 0.129% were false positives (*P*_migration_ < *P*_isolation_ and robustness ≥ 0.95) and 0.409% were ambiguous cases for which ABC did not provide any robust conclusion (robustness < 0.95). Among the 58,000 simulated datasets in which current isolation was assumed (SI, AM; Fig. 2-B), 99.649% were true positives (*P*_isolation_ > *P*_migration_ and robustness ≥ 0.95), 0.002% were false positives (*P*_isolation_ < *P*_migration_ and robustness ≥ 0.95) and 0.34% were ambiguous cases (robustness < 0.95). When current gene flow was assumed, the rates of false positive and ambiguity were very low at every level of population divergence. When current isolation was assumed, a higher rate of ambiguity, but no elevation of the rate of false inference, was observed at low levels of divergence (*Da* < 0.01, Fig. 2-D). This contrasts with the recent suggestion that the full-likelihood method developed in the IMa2 software [32] might be biased towards supporting current gene flow when isolation is recent [19,33] - our approach appears to be immune from this bias. To specifically address this point, we repeated the exact same simulations as in [33] and confirmed that our ABC approach has a reduced power (i.e., more ambiguous cases with robustness <0.95) when the split is recent but still a very low rate of false positive in these conditions (see text S1).

**Figure 2.**
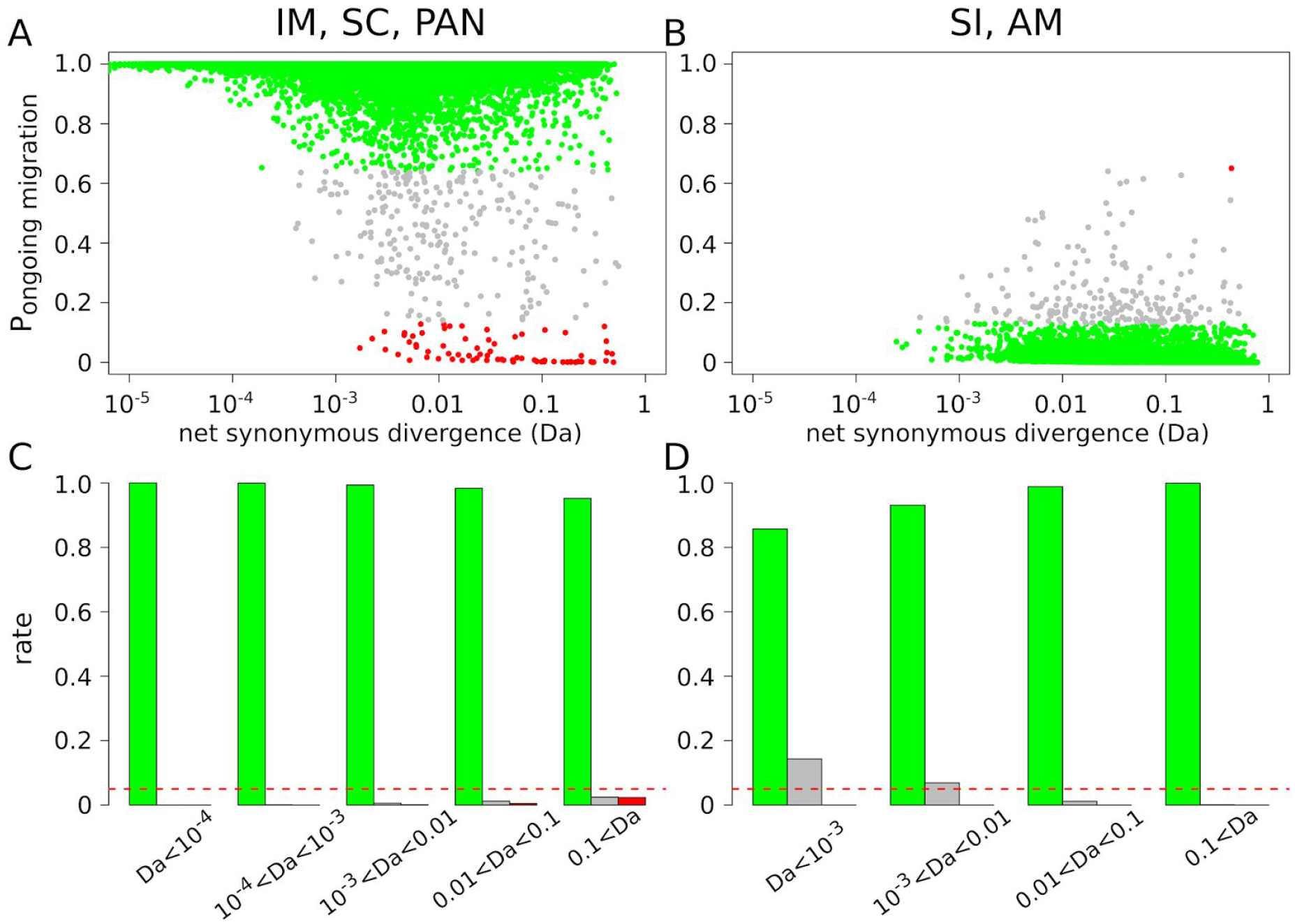
ABC analysis of randomly simulated datasets. Posterior probability *P*_migration_ to support ongoing migration was estimated for a total of 116,000 simulated datasets across 16 models. A. *P*_migration_ as a function of the net synonymous divergence *Da*. Dots represent datasets simulated under the IM, SC and PAN models. The colors show datasets for which gene flow is correctly supported (green) or wrongly rejected (red). Grey dots represent datasets for which the robustness of the ABC analysis is <0.95. B. *P*_migration_ as a function of the net synonymous divergence *Da*. Dots represent datasets simulated under the SI or AM models. The colors show datasets for which gene flow is correctly rejected (green; robustness ≥ 0.95) or wrongly supported (red; robustness ≥ 0.95). C. Proportion of true positives (green), false positives (red) and ambiguous analyses (grey) for different ranges of *Da* across IM, SC and PAN datasets. Horizontal red line shows 5%. D. Proportion of true positives (green), false positives (red) and ambiguous analyses (grey) for different ranges of *Da* across SI and AM datasets.

In addition, the robustness of the ABC inference was only weakly dependent on the sample size when the number of loci was greater than 100: similar ABC results were indeed obtained when we simulated samples of size two, three, 25 or 50 diploid individuals (Fig. S1). Finally, and importantly, simulations showed that ABC is not accurate enough to discriminate between the IM and SC models. Datasets simulated under SC were assigned to SC with high confidence only when the period of isolation before secondary contact was a large enough proportion of the total divergence time (Fig. S2-A). When relatively short periods of isolation were simulated, the method either assigned the datasets to IM or did not provide an elevated posterior probability to any demographic model (Fig. S2-B).

### Dataset: molecular divergence and population differentiation in 61 taxa

The posterior probability of ongoing gene flow was then estimated in 61 pairs of sampled species/populations (Table S1) showing variable levels of molecular divergence (Table S2). 50 pairs were taken from a recent transcriptome-based population genomic study in animals [30], two individuals per population/species being analysed here. The datasets for the other 11 pairs of animal species were downloaded from the NCBI (Table S1). They correspond to sequences from published studies using either ABC, IMa [32] or MIMAR [34], for which three to 78 diploid individuals were analysed.

To represent the relation between the molecular distance between two lineages and the occurrence of ongoing gene flow in natural populations we computed various measures of divergence: *Da* (relative average divergence corrected for within species diversity: [35]), *Dxy* (absolute average divergence) and F_ST_. In our dataset, *Da* ranged from 5.10^−5^ (French vs. Danish populations of *Ostrea edulis*) to 0.309 (*Crepidula fornicata* vs. *C. plana*) (Fig. S3). As expected, *Da* was strongly correlated to F_ST_, a classical measure of population differentiation, which took values ranging from 0 (between *Anas crecca shemya* and *A. crecca attu*) to 0.95 (between *Camponotus ligniperdus* and *C. aethiops;* Fig. S3-A) and less well to the absolute divergence *Dxy* (Fig. S3-B). The across-loci variance in F_ST_ was minimal for low and high values of *Da* (Fig. S3-B), which reflects an F_ST_ homogeneously low at early stages of divergence, homogeneously high at late stages of divergence, and heterogeneous among genes at intermediate levels of *Da* (Fig. S3).

### Statistical analysis: assessment of ongoing gene flow

For the 61 studied pairs of populations/species, we focused on synonymous positions and investigated the prevalence of ongoing gene flow by estimating the posterior probabilities of 16 different models under ABC. These 16 models represent the combinations of five demographic models (SI, AM, IM, SC and panmixia) and four assumptions regarding the genomic heterogeneity in introgression (for AM, IM and SC only) and drift rates (for all models; see above and Material and Methods). The posterior probability P_migration_ that the two populations currently exchange migrants was estimated by summing the contributions of the PAN, IM and SC models (Fig. 1) and plotted against measures of molecular divergence (Fig. 3). *Da*, which can be understood as the per-site amount of neutral derived mutations being fixed in the different lineages, provided the best relationship (Fig. 3). Results with other measures of divergence and with the estimated age of the split (*T*_*split*_ parameter under the IM model) are also shown (Fig. S4-S7).

**Figure 3.**
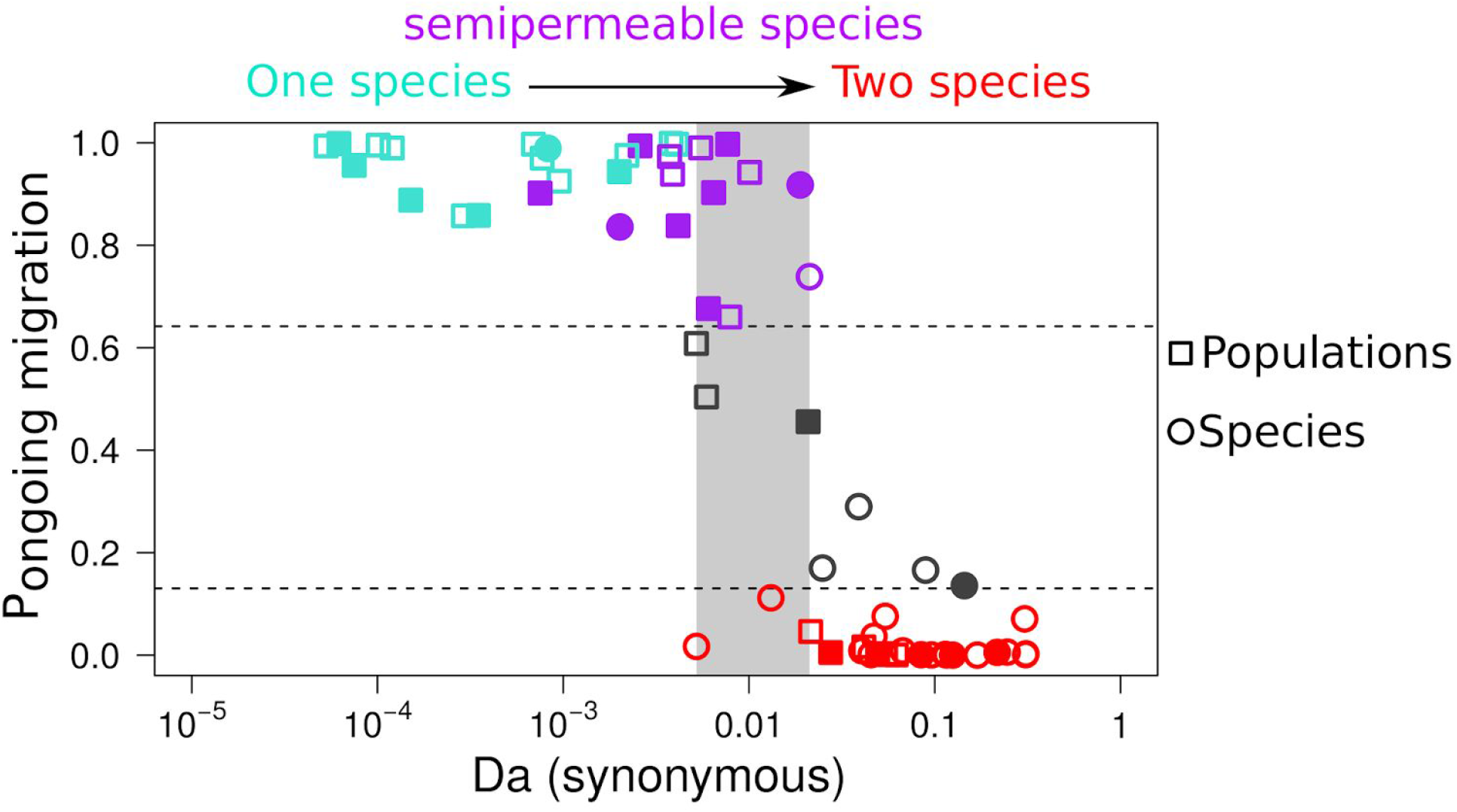
Probability of ongoing gene flow along a continuum of molecular divergence. Each dot is for one observed pair of populations/species. X-axis: net molecular divergence *Da* measured at synonymous positions (log10 scale) and averaged across sequenced loci. Y-axis: relative posterior probability of ongoing gene flow (i.e., SC, IM and PAN models) estimated by ABC. Red dots: pairs with a strong support for current isolation. Grey dots: with no strong statistical support for any demographic models (robustness <0.95). Blue dots: pairs with strong statistical support for genome-homogeneous ongoing gene flow. Purple dots: pairs with strong statistical support for genome-heterogeneous ongoing gene flow. Filled symbols: pairs with a strong support for genome-heterogeneous *Ne*. Open symbols: genomic-homogeneous *Ne*. The light grey rectangle spans the range of net synonymous divergence in which both currently isolated and currently connected pairs are found.

Over the continuum of divergence, the 22 pairs with *Da* lower than 0.5% received a support for ongoing gene flow with a robustness ≥0.95 (Fig. 3). The first identified semi-permeable barrier to gene flow was detected at *Da*≈0.075%, a pair of *Malurus* (fairywren) species [36] for which ABC strongly supports heterogeneity in *M*. When the net divergence was between 0.5% and 2%, inferences about gene flow were variable, and sometimes uncertain. In this grey zone, gene flow was strongly supported in seven pairs, always with a strong support for genomic heterogeneity in introgression rates. Still in the grey zone, ABC did not distinguish between isolation and introgression in three pairs of species and provided strong support for isolation in two other pairs. Finally, among the 27 most divergent pairs of species where *Da* was greater than 2%, we found 23 pairs with a strong support for current isolation and four ambiguous pairs (Fig. 3).

We investigated the impact of assumptions about genomic heterogeneity in *Ne* and *M* on the detection of current introgression (Fig. S4–S9). When both parameters were allowed to vary among loci, pairs of populations with *Da* exceeding 0.1% and showing strong statistical support for ongoing migration tended to obtain support for genomic heterogeneity in introgression rates. But when constant introgression rate was assumed (homoM_heteroN and homoM_homoN models), the importance of gene flow became underestimated in several divergent pairs of species, consistent with previous reports (e.g. [15]). When we compared models assuming homogeneous *versus* heterogeneous effective population size across loci, we found that the former tended to overestimate the prevalence of ongoing gene flow (Fig. S8), again in line with published analyses [19]. Analyses assuming homogeneous *Ne* and *M* in many cases failed to support either isolation or migration, producing the highest number of ambiguous pairs (Fig S8). The detected genomic heterogeneity in gene flow increased with *Da* until 2% of divergence. Finally, across the whole continuum, there was no significant effect of the divergence on the probability of supporting genomic heterogeneity in effective population size in our data set.

### No effect of habitat, geography, phylogeny or life-history traits

We investigated the influence of a number of ecological, geographical, phylogenetic and life-history variables on the posterior probability of ongoing gene flow. This was achieved under the heteroM_heteroN model using data from [30]. We detected no significant effect of species longevity or log-transformed propagule size (size of the developmental stage that leaves the mother and disperses) on the log-transformed probability of ongoing gene flow. In the same vein, marine organisms (*n*=25) did not exhibit a higher propensity for ongoing gene flow than terrestrial ones (*n*=36; r^2^ below 0.01%). The log-transformed probability of ongoing gene flow was significantly higher (*p*-val=0.002, r^2^=0.14) in vertebrates (*n*=20) than in invertebrates (*n*=41), but the effect disappeared when the level of divergence was controlled for (net synonymous divergence<0.04: 17 vertebrate pairs, 22 invertebrate pairs, *p*=0.32, r^2^=0.03). This effect only reflects the paucity of pairs of vertebrate population/species with a high divergence in our data set. Finally, we tested whether the current geographic distribution of species coincides with the establishment of genetic structure in our data by distinguishing pairs in which the two considered species/populations occur on the same *vs*. distinct continents or oceans. We did not find any significant effect of this variable on the estimated values of *P*_migration_ in either of the three divergence zones: *Da*<0.5%, t-test = −0.015269, df = 18.522, *P* value = 0.988; 0.5%<*Da*<2%, t-test = −0.74229, df = 7.1996, *P* value = 0.4814; 2%<*Da*, t-test =0.35512, df = 22.426, *P* value = 0.7258.

### Ongoing gene flow and taxonomic status

Finally, we verified whether our inferences confirmed or contradicted the current taxonomy (table S3). Our dataset comprises 26 pairs of recognized species and 35 pairs of recognized populations, or sub-species, sharing a common binomen. Twenty-one pairs of recognized species belonged to the high-divergence zone (*Da*>0.02). Of these, 16 were inferred to be currently isolated, four produced ambiguous results and one pair, *Eunicella cavolinii* vs. *E. verrucosa* (gorgonian), was found to be connected by heterogeneous gene flow. Among the five remaining recognized pairs of species (with *Da*<0.02), two were inferred as being fully isolated and three were inferred to be connected species: two pairs of semi-isolated species with heterogeneous gene flow (*Mytilus galloprovincialis* vs *M. edulis* and *Macaca mulatta* vs *M. fascicularis*) and the *Gorilla gorilla* vs. *G. beringei* pair, which was found to be connected by homogeneous gene flow. Of the 35 pairs of recognized populations from the same species, 6 with *Da*>0.02 were inferred to be isolated - cryptic species. Genetic isolation has been previously suspected between northern and southern populations of *Pectinaria koreni* (trumpet worms) [37], between the blue and purple morphs of *Cystodytes dellechiajei* (colonial ascidians) [38], and between the L1 and L2 lineages of *Allolobophora chlorotica* (earthworms) [39], but is here newly revealed between Morrocan and European populations of *Melitaea cinxia* (Glanville fritillary), between Spanish and French populations of *A. chlorotica* L2 and between Mediterranean and tropical populations of *Culex pipiens*.

## Discussion

We performed a comparative speciation genomics analysis in 61 pairs of populations/species from various phyla of animals. Our ABC analysis, which takes into account the confounding effect of linked selection heterogeneity, provides a first global picture of the prevalence of gene flow between diverging gene pools during the transition from one to two species.

### Accounting for among-loci heterogeneity in drift and migration rate

Inferring the history of divergence and gene flow, which determines the rate of accumulation of species barriers, is of prime importance to understand the process of speciation [17]. This can be achieved by various methods, among which ABC approaches have proven particularly flexible and helpful to compare alternative evolutionary models. Our analysis of simulated datasets illustrates that ABC methods have the power to effectively discriminate recent introgression *versus* current isolation based on datasets of several hundreds of loci and a few individuals per species - typical of population genomic studies. Comparisons of alternative demographic models, however, can be strongly impacted by assumptions regarding the genomic distribution of effective population size (*Ne*) and introgression rate (*M*). Heterogeneities in *Ne* and *M* are common in natural populations as a result of selective processes applying either globally (background selection [19,40,41]) or specifically against migrants (genetic barriers [12, 42]).

Following [13], we here introduced a framework in which each of the two effects, or both, can be readily accounted for. In our analysis, the number of pairs of populations/species for which ambiguous conclusions were reached was maximal when genomic heterogeneities of both migration and drift were neglected. Incorporating within genome variation in *Ne* tended to enhance the support for models with current isolation, as previously suggested [19]. The heteroN model makes a difference regarding inference of current gene flow between the highly divergent *Ciona intestinalis* and *C. robusta* species (see below). Conversely, incorporating heterogeneity in *M* doubled the number of pairs for which ongoing gene flow was supported, when compared to analyses with homogenous *M* where most of these pairs exhibited ambiguous results. Our study therefore underlines the importance of accounting for genomic heterogeneities for both *Ne* and *M* when comparing alternative models of speciation [14,15,19], and calls for prudence regarding the conclusions to be drawn from the analysis of a single pair. However, it is important to recall here that the action of natural selection on its molecular target and neighborhood is more complex than a simple reduction in *Ne*. Our modeling of genomic heterogeneity in drift and selection by a Beta distribution of *Ne* throughout genome is an approximation which can not replace an explicit modeling of these processes. In our modeling, we assumed that a given locus *i* is independently affected by drift and selection in all of the simulated populations including the ancestral. Our choice was motivated by the generality of this model. An alternative approach to model genomic heterogeneity in *Ne* can be assuming that background selection is the main process shaping genomic landscapes of diversity. This can be approximated by assuming that a locus *i* is equally affected by drift and selection in all populations instead of assuming independent effects as in our study.

Among models assuming ongoing gene flow, our ABC analysis of simulated and empirical data often failed to discriminate between the Isolation-with-Migration and Secondary Contact models. These two models yield similar signatures in genetic data, so that only relatively recent secondary contacts following long periods of interrupted gene flow can be detected with high confidence (Fig. S2-D) [43]. Similarly, among models excluding ongoing gene flow, distinguishing between Strict Isolation and Ancient Migration was not possible in a substantial number of cases. These are challenges for future methodological research in the field, with important implications regarding the debate about the requirement of geographic isolation to complete speciation [7,44]. Only two diploid individuals per population/species were used in this analysis, for the sake of comparability between data sets (in many populations no more than two individuals are available), and due to computational limitation. However, our evaluation of the effect of sample size on ABC-based demographic inference suggested that two individuals per population were largely sufficient to capture the main signal when more than 100 loci are available (Fig. S1), which mainly has its sits in coalescent events shared by every individuals of a population sample [45].

### Prevalent gene flow between slightly diverged gene pools

Although ABC analyses of particular pairs of populations can be affected by the choice of model of genomic heterogeneity, the overall relationship between net molecular divergence and detected ongoing gene flow was qualitatively similar among analyses. Pairs of populations diverging by less than 0.5% were found to currently exchange migrants. This includes populations that form a single panmictic gene pool, and pairs of diverging populations/species connected by gene flow. The low-divergence area contains pairs of populations showing conspicuous morphological differences, such as Eastern vs. Western gorilla or the *cuniculus* and *algirus* subspecies of rabbit (*Oryctolagus cuniculus*).

No pair of populations in this range of divergence was supported to be genetically isolated or yielded ambiguous results. Simulations indicate that our ABC approach is not expected to yield false inference of gene flow in recently isolated populations, contrary to what was suggested with the full-likelihood approach of IMa2 [33]. The main risk is rather a false inference of isolation despite gene flow (Fig. 2), which can be explained by the fact that the SI model is less parameterized than models assuming gene flow (IM and SC). ABC had a low false positive rate even when we simulated very recent splits as in previous papers [19,33]. This is probably because, in strict isolation, shared polymorphisms are quickly sorted into private polymorphisms and fixed differences after population split, such that *Da* can hardly be very small in the absence of gene flow [46]. Our analysis therefore identifies *Da*<0.5% as a good synthetic proxy to attest for the existence of gene flow. Other measures of divergence, although producing a qualitatively similar pattern, did not predict the existence of current gene flow as nicely as Da did.

Pairs in the low range of divergence must correspond to populations that did not accumulate sufficiently strong and numerous genetic barriers, so that gene flow currently occurs at important rates. The detection of significantly heterogeneous introgression rates in a number of low-diverged pairs (*Da*<0.5%) demonstrates the ability of our ABC approach to detect semi-permeable barriers quite efficiently at early stages of speciation and supports the rapid evolution of Dobzhansky-Muller incompatibilities [47,48]. A majority of the pairs from the low-divergence area, however, did not yield any evidence for among-loci heterogeneity of introgression rate. Some might correspond to effectively isolated backgrounds that are missed by our method by lack of power when the signal is too tenuous. It is quite plausible, however, that some pairs of populations/species in the low-divergence zone have differentially fixed mutations with major effects on hybrid fitness whereas other have not, due to mutational stochasticity and/or across-taxa differences in the genetic architecture of barriers – i.e., simple (two locus) vs. complex incompatibilities, and strength of associated selective effects [49].

### Suppressed gene flow at high sequence divergence

At the other end of the continuum, it appears that above a divergence of a few percent, barriers are strong enough to completely suppress gene flow: almost all pairs of species with *Da* > 2% were found to have reached reproductive isolation with strong support. This might result from impaired homologous recombination due to improper pairing of dissimilar homologous chromosomes at meiosis, which would reduce the fecundity of hybrids [50,51]. Of note, the upper threshold for reproductive isolation (*Da*=2%, *Dxy*=5.5%) is of the order of magnitude of the maximal level of within-species genetic diversity reported in animals [30,52], somewhat consistent with the hypothesis of a physical constraint imposed by sequence divergence on the ability to reproduce sexually. Alternatively, the 2% figure may represent a threshold above which Dobzhansky-Muller incompatibilities are normally in sufficient number and strength to suppress introgression. The two hypotheses are not mutually exclusive, but pertain to distinctive processes of genetic isolation; the former would be maximally expressed during F1 hybrid meiosis, while the latter would affect recombined, mosaic individuals carrying alleles from the two gene pools at homozygous state.

In the high-divergence area, no instance of among-loci heterogeneous migration was detected, indicating that introgression is blocked across the whole genome in these pairs of species. A number of highly divergent species pairs yielded support for among loci heterogeneous *Ne*, suggesting that the same regions of the genome are under strong background selection in the two diverging entities – presumably regions of reduced recombination and/or high density in functional elements. Neglecting the genomic heterogeneity in *Ne* can lead to false inference of gene flow. For instance, allowing genomic heterogeneity in *M* but not in *Ne* led to strong statistical support for a secondary contact between the highly divergent *Ciona intestinalis* (formerly *C. intestinalis B*) and *C. robusta* (formerly *C. intestinalis* A) species (Fig. S4-S5), consistent with [14], but accounting for heterogeneity in both *M* and *Ne* resulted in an ambiguous result without a sufficiently strong support for any models. The among-loci variance in differentiation between these two species, which was interpreted as mainly reflecting introgression at few loci in [14], is shown here to possibly be the result of a more complex situation that our models failed to capture.

### Intermediate divergence levels: the grey zone of speciation

The area of intermediate divergence from 0.5% to 2% of net synonymous divergence unveils the grey zone of the speciation continuum. In this grey zone, isolated pairs of populations/species coexist with pairs connected by migration, and the latter are mainly composed of semi-isolated genetic backgrounds, the situation under which taxonomic conundrums flourish. Cases of ambiguous conclusions about the demographic history also tended to be found in this intermediate zone, perhaps reflecting instances of complex divergence models that are not well predicted by our demographic models. Researchers should be ready to face problems regarding demographic inference, and therefore parameter estimation, when conducting a project of speciation genomics falling in the grey zone. Accounting for genomic heterogeneity of introgression and drift rates appears to be crucial for detecting current gene flow in this range of divergence (Figures S4-S7). For instance, the mussel species *M. galloprovincialis* vs *M. edulis* and the gorgonian species *Eunicella cavolinii* vs *E. verrucosa* are the two most divergent pairs for which ongoing introgression was detected, but this only appeared when the genomic variation in *M* was accounted for – the homoMhomoN and homoMheteroN models yielded ambiguous conclusions about these pairs of species, in which the existence of semi-permeable barriers has previously been demonstrated [53,54].

Our analysis revealed significant among-loci heterogeneous migration in as many as thirteen pairs of populations/species (Fig. 3). This illustrates the commonness of semi-permeable genomes at intermediate levels of speciation, when some, but not all, genomic regions are affected by barriers to gene flow. Besides mussels and gorgonians, heterogeneous gene flow was newly detected between American and European populations of *Armadillidium vulgare* (wood lice) and *Artemia franciscana* (brine shrimp), between Atlantic and Mediterranean populations of *Sepia officinalis* (cuttlefish), and between the closely related *Eudyptes chrysolophus moseleyi* vs. *E. c. filholi* (penguins) and *Macaca mulatta* vs. *M. fascicularis* (macaques) – in addition to the previously documented mouse [55], rabbit [56] and fairywren [57] cases.

The grey zone, finally, includes populations between which unsuspected genetic isolation was here revealed, such as the Moroccan vs. European populations of *Melitaea cinxia* (Glanville fritillary), and the Spanish vs. French populations of *A. chlorotica* L2 (earthworm), which according to our analysis correspond to cryptic species. Our genome-wide approach and proper modeling of heterogeneous processes therefore clarified the status of a number of pairs from the grey zone, emphasizing the variety of situations and the conceptual difficulty with species delineation in this range of divergence.

### Implications for speciation and conservation research

Our dataset is composed of a large variety of taxa with deep phylogenetic relationships and diverse life history traits. In principle, the propensity to evolve pre-zygotic barriers might differ between groups of organisms (e.g. broadcast spawners versus copulating species, [58]). We did not detect any significant effect of species biological/ecological features or taxonomy on the observed pattern. Highly polymorphic broadcast spawners and low diversity large vertebrates with strong parental investment were equally likely to undergo current gene flow, for a given divergence level. Whether the pace of accumulation of genetic barriers, the so-called speciation clock, varies among taxonomic group is a major challenge in speciation research and requires the dissection of the temporal establishment of barriers in many different taxas [59,60]. State-of-the-art ABC methods offer the opportunity to investigate the genome-wide effect of barriers to gene flow in natural populations but cannot provide answers about how and why barriers have evolved. However, our report of a strong and general relationship between molecular divergence and genetic isolation across a wide diversity of animals suggests that, at the genome level, speciation operates in a more or less similar fashion in distinct taxa, irrespective of biological and ecological peculiarities.

Interestingly, we did not detect any significant effect of geographic range overlap. This result may appear as unexpected at first sight, because one expects gene flow to be dependent on geography. One explanation could be that we used too crude a measure of range overlap. Alternatively, this result could support the idea that in many taxa the observed genetic structure was established in the past in a geographic context different from the current one, and only recently reshuffled by recent migration/colonization processes [61]. According to this hypothesis, genetic subdivision could have little to do with contemporary connectivity.

The width of the grey zone indicates that a number of existing taxonomic debates regarding species definition and delineation are difficult by nature and unlikely to be resolved through the analysis of a limited number of loci. Most of the molecular ecology literature, however, is based on datasets consisting of mitochondrial DNA and rarely more than a dozen microsatellite loci. The time when genome-wide data will be available in most species of interest is approaching though not yet reached. Since then, we have to accept that knowledge about the existence of gene flow between diverged entities could not be settled from genetic data alone in a substantial fraction of taxa. In addition, our study highlights the commonness of semi-isolated entities, between which gene flow can be demonstrated but only concerns a fraction of loci, further challenging the species concept. We should therefore be prepared to make decisions regarding conservation and management of biodiversity in absence of well-defined species boundaries.

## Materials and Methods

### Taxon sampling

A total of 61 pairs of populations/species of animals were analyzed (Table S1). These include 10 pairs taken from the speciation literature and 51 pairs newly created here based on a recently published RNAseq dataset [30], which includes 96 species of animals from 31 distinct families and eight phyla, and one to eleven individuals per species. Twenty-nine of the newly created pairs corresponded to distinct populations within a named species. Populations were here defined based on a combination of geographic, ecotypic and genetic criteria: we contrasted groups of individuals (i) living in allopatry and/or differing in terms of their ecology, and (ii) clustering as distinct lineages in a neighbour-joining analysis of genetic distances between individuals. The two most covered individuals per population were selected for ABC analysis. In four species three distinct populations were identified, in which case the three possible pairwise comparisons were performed. Results were qualitatively unchanged when we kept a single pair per species. Twenty-two of the newly created pairs consisted of individuals from two distinct named species that belonged to the same family. Again, the two most covered individuals per species were selected for analysis. In the case of species in which several populations had been identified, we chose to sample two individuals from the same population for between-species comparison. When more than two species from the same family were available, we selected a single pair based on a combination of sequencing coverage and genetic distance criteria, comparisons between closely related species being favored. Raw and final datasets are available from the PopPhyl website (http://kimura.univ-montp2.fr/PopPhyl/). Sample sizes, number of loci and source of data are listed in Table S1.

### Transcriptome assembly, read mapping, coding sequence prediction

For the 51 recently obtained pairs, Illumina reads were mapped to predicted cDNAs (contigs) with the BWA program [62]. Contigs with a per-individual average coverage below ×2.5 were discarded. Open reading frames (ORFs) were predicted with the Trinity package [63]. Contigs carrying no ORF longer than 200 bp were discarded. In contigs including ORFs longer than 200 bp, 5′ and 3′ flanking non-coding sequences were deleted, thus producing predicted coding sequences that are hereafter referred to as loci.

### Calling single nucleotide polymorphisms (SNPs) and genotypes

At each position of each locus and for each individual, diploid genotypes were called using the reads2snps program [64]. This method first estimates the sequencing error rate in the maximum-likelihood framework, calculates the posterior probability of each possible genotype, and retains genotypes supported at >95% if ten reads per position and per individual were detected. Possible hidden paralogs (duplicated genes) were filtered using a likelihood ratio test based on explicit modeling of paralogy. For our demographic inferences only synonymous positions were retained. Synonymous length and positions were then computed for each loci using polydNdS [65].

### Summary statistics

For all of the 61 pairs of populations/species, we calculated an array of 31 statistics widely used for demographic inferences [31,34,66,67]. The average and standard variation over loci for: (1) the number of biallelic positions; (2) the number of fixed differences between the two gene pools; (3) the number of polymorphic sites specific to each gene pools; (4) the number of polymorphic sites existing in both gene pools; (5) Wald and Wolfowitz statistics [68]; (6) Tajima's pi [69]; (7) Watterson's theta [70]; Tajima's *D* for each gene pools [71]; (8) the gross divergence between the two gene pools (*Dxy*); (9) the net divergence between the two gene pools (*Da*); (10) F_ST_ measured by 1-p_w_/p_T_ where p_W_ is the average allelic diversity based on the two gene pools and p_T_ is the total allelic diversity over the two gene pools; (11) the Pearson's R^2^ correlation coefficient in p calculated between the two gene pools. Observed values of summary statistics are summarized for each species in table-S2.

### Demographic models

Five distinct demographic models were considered: panmixia (PAN), Strict Isolation (SI), Ancestral Migration (AM), Isolation with Migration (IM) and Secondary Contact (SC, Fig. 1). The PAN model assumes that the two investigated gene pools are sampled from a single panmictic population of size *Ne*. The SI model describes the subdivision of an ancestral panmictic population of size *N*_*anc*_ in two isolated gene pools of sizes *N*_*pop-1*_ and *N*_*pop-2*_. The two sister gene pools then evolve in absence of gene flow. Under the IM model, the two sister gene pools that split *T*_*spli*t_ generations ago continuously exchange alleles as they diverge. Under the AM model gene flow occurs between *T*_*spli*t_ and a more recent *T*_*AM*_ date, after which the two gene pools evolve in strict isolation. The SC model assumes an early divergence in strict isolation followed by a period of gene flow that started *T*_*SC*_ generations ago.

### Heterogeneity in introgression and effective population size

We assumed that the effects of selection on linked sites can be described in terms of heterogeneous effective population size (putatively affecting all demographic models) and/or migration rate (only affecting the IM, AM and SC models). In the homoM setting, one gene flow parameter (*M=N.m*) is randomly sampled from a uniform prior distribution for each direction. *M*_*1*_ is the direction from gene pool 2 to gene pool 1 and *M*_*2*_ is the direction from gene pool 1 to gene pool 2. All loci share the same *M*_*1*_ and *M*_*2*_ values, but *M*_*1*_ and *M*_*2*_ are independently sampled. In the heteroM setting a specific migration rate is attributed per locus and per direction of migration. Thus, for each direction, a hyper-prior is first randomly designed as a Beta distribution. A value of *M*_*1,i*_, and *M*_*2,i*_ is then drawn for each loci *i* from the two hyper-priors. In the homoN setting, the effective population sizes *N*_*anc*_ (ancestral population), *N*_*pop-1*_ (gene pool 2) and *N*_*pop-2*_ (gene pool 2) are independent but shared by all loci. In the heteroN setting, heterogeneity in effective population size is independently modeled for the three populations (ancestor, gene pool 1 and gene pool 2). For each population, a proportion a of loci is assumed to evolve neutrally and share a common value for *N*_*anc*_, *N*_*pop-1*_ or *N*_*pop-2*_, *a* being sampled from the uniform prior [0 - 1]. The remaining loci, in proportion 1-*a*, are assumed to be affected by natural selection at linked loci. They are assigned independent values of *N*, which are sampled from Beta distributions defined on the intervals [0 - *N*_*anc*_], [0 - *N*_*pop-1*_] and [0 - *N*_*pop-2*_]. In this setting a and *Ne* differ between the three populations but are sampled from distributions sharing the same shape parameters.

### Approximate Bayesian Computation

The combination of demographic models and genomic settings resulted in a total of 16 distinct models, namely the homoN and heteroN versions of PAN and SI, and the homoM_homoN, homoM_heteroN, heteroM_homoN, heteroM_heteroN versions of IM, AM and SC. Model fit assessment and parameter estimation were performed under the ABC framework. 3,000,000 of multilocus simulations under each model were conducted using the coalescent simulator msnsam [66,72]. For each of the 61 pairs of populations/species the posterior probability of each model was estimated using a feed-forward neural network implementing a nonlinear multivariate regression by considering the model itself as an additional parameter to be inferred under the ABC framework using the R package “abc” [73]. The 10,000 replicate simulations (out of 16 x 3,000,000) falling nearest to the observed values of summary statistics were selected, and these were weighted by an Epanechnikov kernel that peaks when *S*_obs_ = *S*_sim_. Computations were performed using 50 trained neural networks and 10 hidden networks in the regression. The posterior probability of each model was obtained by averaging over ten replicated ABC analysis.

### Robustness

Among a set of compared models, ABC returns a best-supported model *M* and its posterior probability *P*_*M*_. The returned model is validated when *P*_*M*_ is above an arbitrary threshold *X*, corresponding to the posterior probability above which the statistical support for a model is considered as being significant. The robustness of the inference, *i.e.*, the probability to correctly support model *M* if true, obviously depends on *X*. To assess the reliability of our approach, we randomly simulated 116,000 pseudo-observed datasets (PODs) distributed over the 16 compared models. Simulations were independent of the 3,000,000 x 16 reference simulations used for model comparisons in our main analysis, but their parameters share the same boundaries.

For each simulated POD, we estimated the posterior probabilities *P*_*i*_ of the 16 compared models through ABC. The probability of correctly supporting *M* given *X* was calculated as: 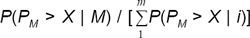, where 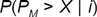 is the probability that a dataset simulated under *m* will be supported by ABC as being *M* with a posterior probability above *X* [31]. This is the proportion, among simulated datasets inferred by ABC to correspond to *M*, of those actually generated under *M*.

For the “ongoing gene flow” *vs* “current isolation” model comparison, we empirically measured that robustness to support gene flow starts to be above 0.95 if *P*_migration_ ≥ 0.6419 and the robustness to support isolation is above 0.95 if *P*_migration_ ≤ 0.1304. For datasets with *P*_migration_ between 0.1304 and 0.6419, we did not attribute a best model but treated them as “ambiguous cases”.

All of the informatic codes and command lines used to produce the analysis are openly available online (https://github.com/popgenomics/popPhylABC).

## Acknowledgments

We thank Aude Darracq, Vincent Castric, Pierre-Alexandre Gagnaire, Xavier Vekemans and John Welch for insightful discussions. The computations were performed at the Vital-IT (http://www.vital-it.ch) Center for high-performance computing of the SIB Swiss Institute of Bioinformatics and the ISEM computing cluster at the platform Montpellier Bioinformatique et Biodiversité of the LabEx CeMEB. NB was funded by the Agence Nationale de la Recherche (HYSEA project, ANR-12-BSV7- 0011). This is article 2016-XXX of Institut des Sciences de l’Evolution de Montpellier.

---

**Figure S1.**
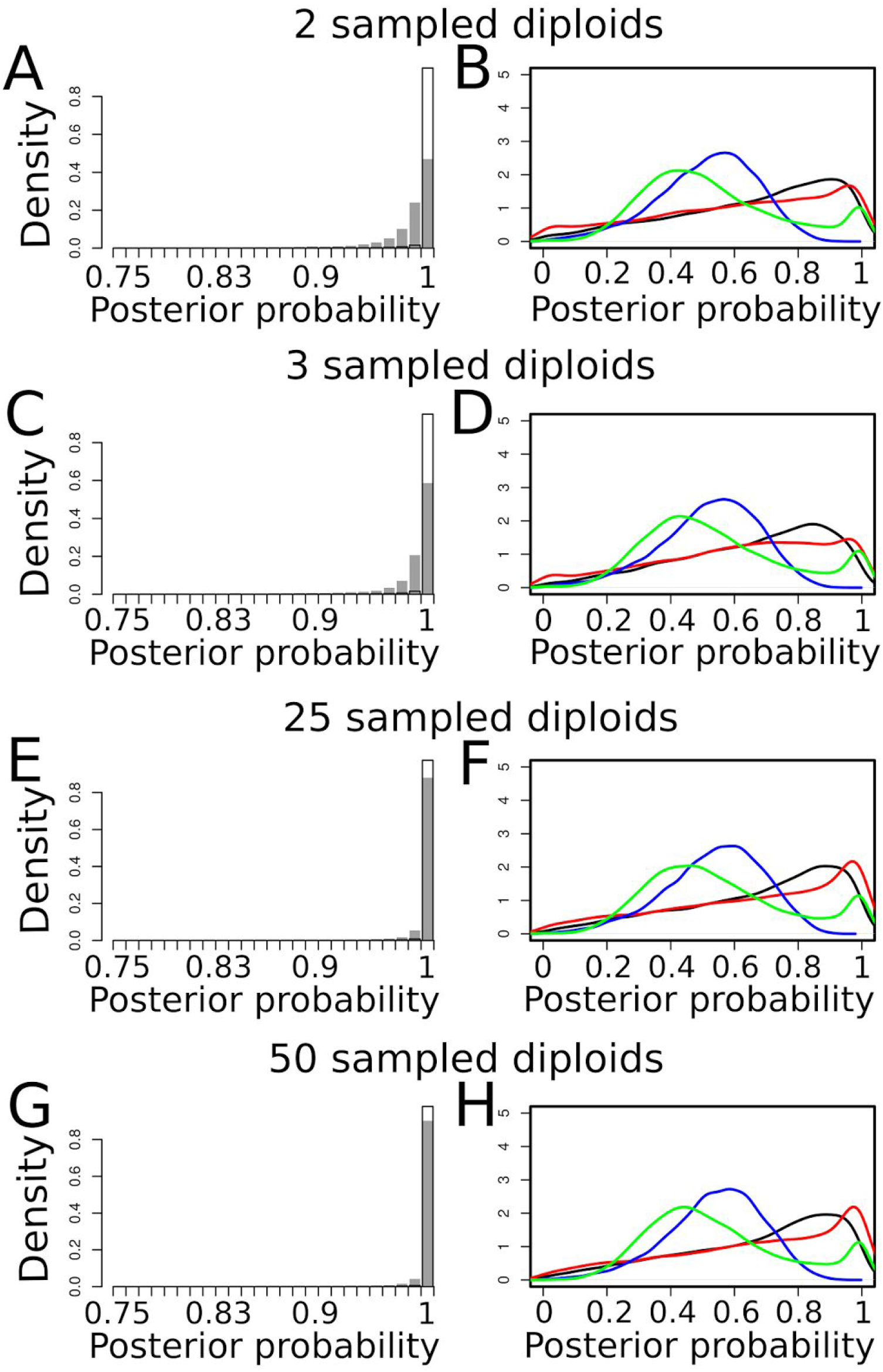
Effects of the number of sampled individuals on robustness of model comparisons when 100 loci are investigated. Analyses were made by simulating four different datasets: A-B: 100 loci sampled in two diploid individuals in each daughter species. C-D: 100 loci sampled in three diploid individuals in each daughter species. E-F: 100 loci sampled in 25 diploid individuals in each daughter species. G-H: 100 loci sampled in 50 diploid individuals in each daughter species. Panels on the left border show the distributions of *P*(current isolation | current isolation) (white bars) and *P*(current introgression | current introgression) (grey bars) measured after ABC analysis of 20,000 PODs simulated under each models. Panels on the right border show the distributions of *P*(SI | SI) (black lines), *P*(AM | AM) (red lines), *P*(IM | IM) (blue lines) and *P*(SC | SC) (green bars) measured after ABC analysis of 20,000 PODs simulated under each models.

**Figure S2.**
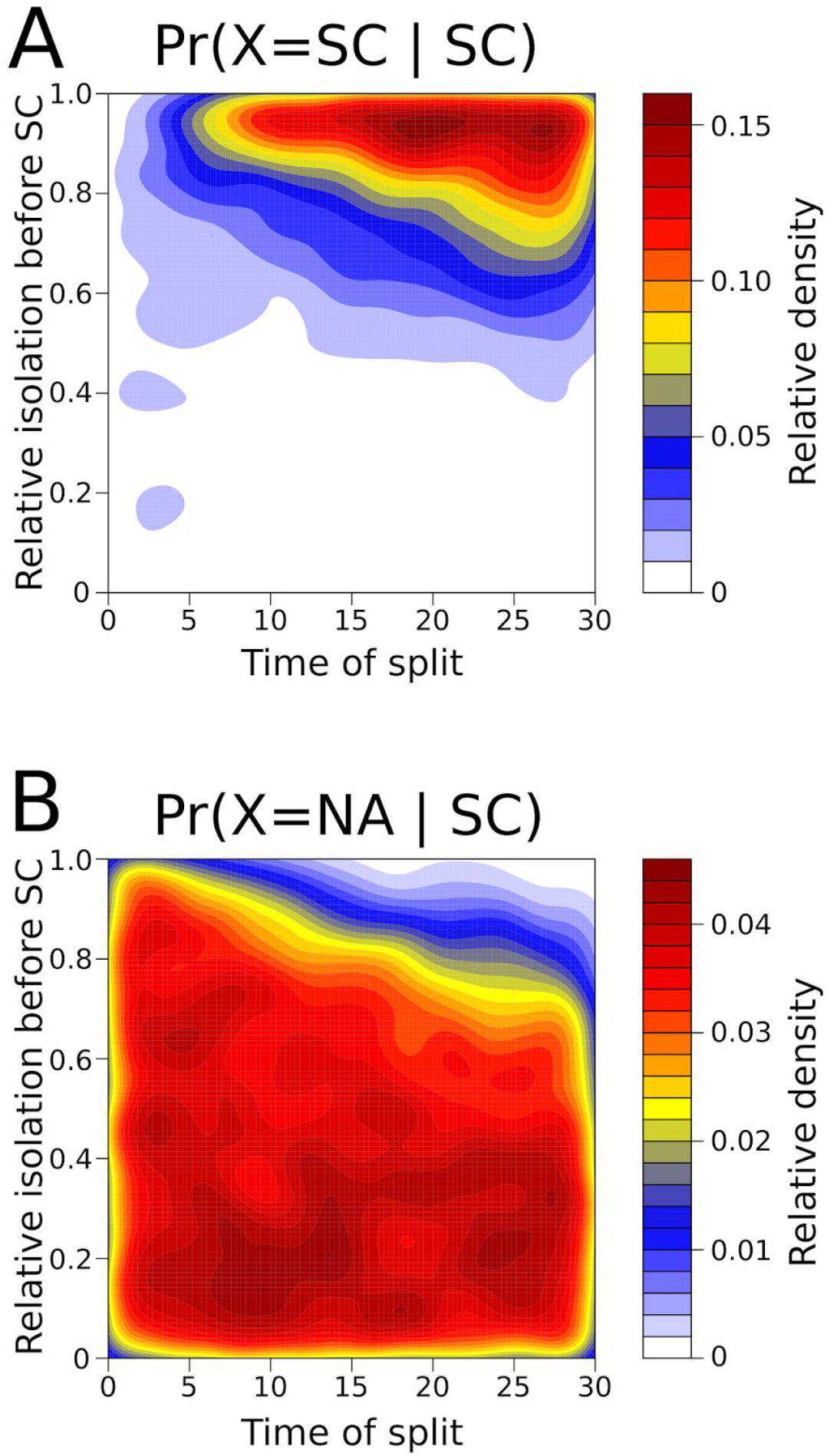
Effect of parameter combinations on the correct support of the SC model. A. Two-dimensional space of parameters of the SC model showing simulations leading to a correct support of SC (i.e *P*(SC | SC) > 0.8). X-axis represents the time since the ancestral split. Y-axis represents the relative time the two daughter species remained isolated before the secondary contact. Colors represent the density in simulations with *P*(SC | SC) > 0.8. B. Two-dimensional space of parameters of the SC model showing simulations leading to the absence of a robust conclusion using ABC. Colors represent the density in simulations with *P*(NA | SC).

**Figure S3.**
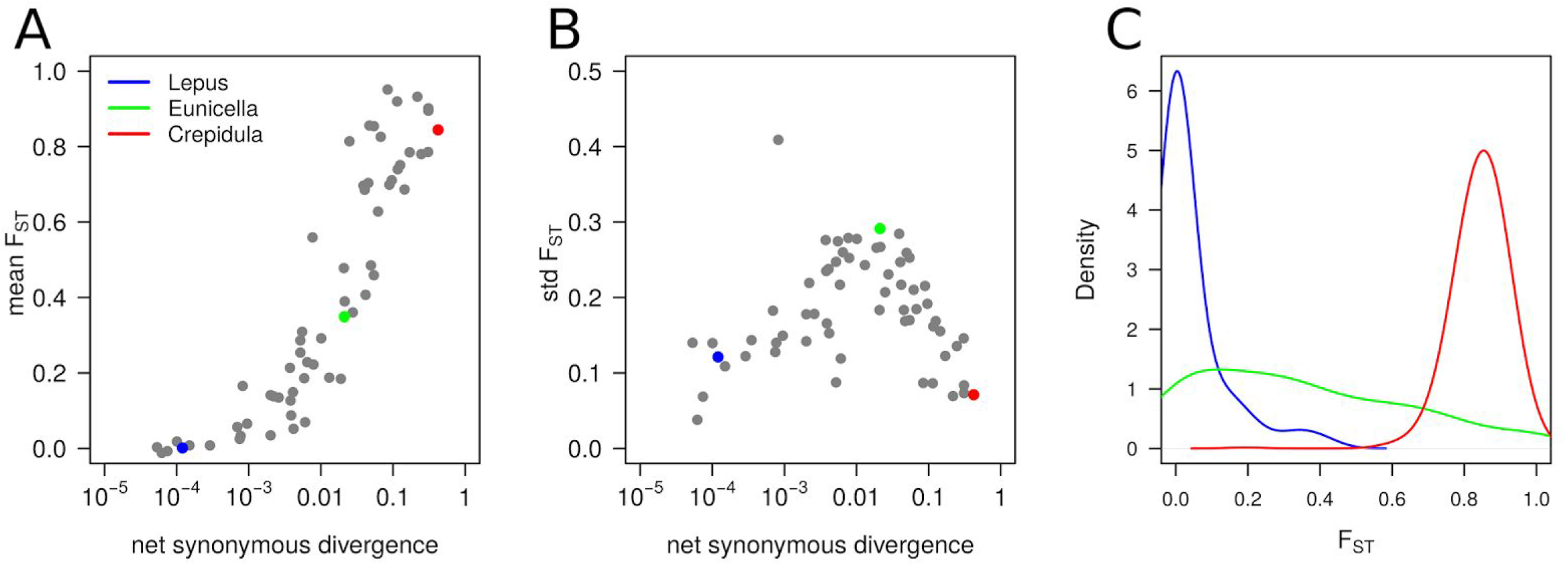
Relation between synonymous divergence and genetic differentiation. A. Each grey dot represents a pair of species/populations. Lepus (Spanish and Portuguese populations of *Lepus granatensis*), Eunicella (*Eunicella cavolinii* and *E. verrucosa*) and Crepidula (*Crepidula fornicata* and *Bostrycapulus aculeatus*) indicate representative pairs of poorly, intermediately and highly divergent species/populations. B. Effect of divergence on across-loci variance in F_ST_. C. Genomic distribution of F_ST_ for the Lepus, Eunicella and Crepidula datasets.

**Figure S4.**
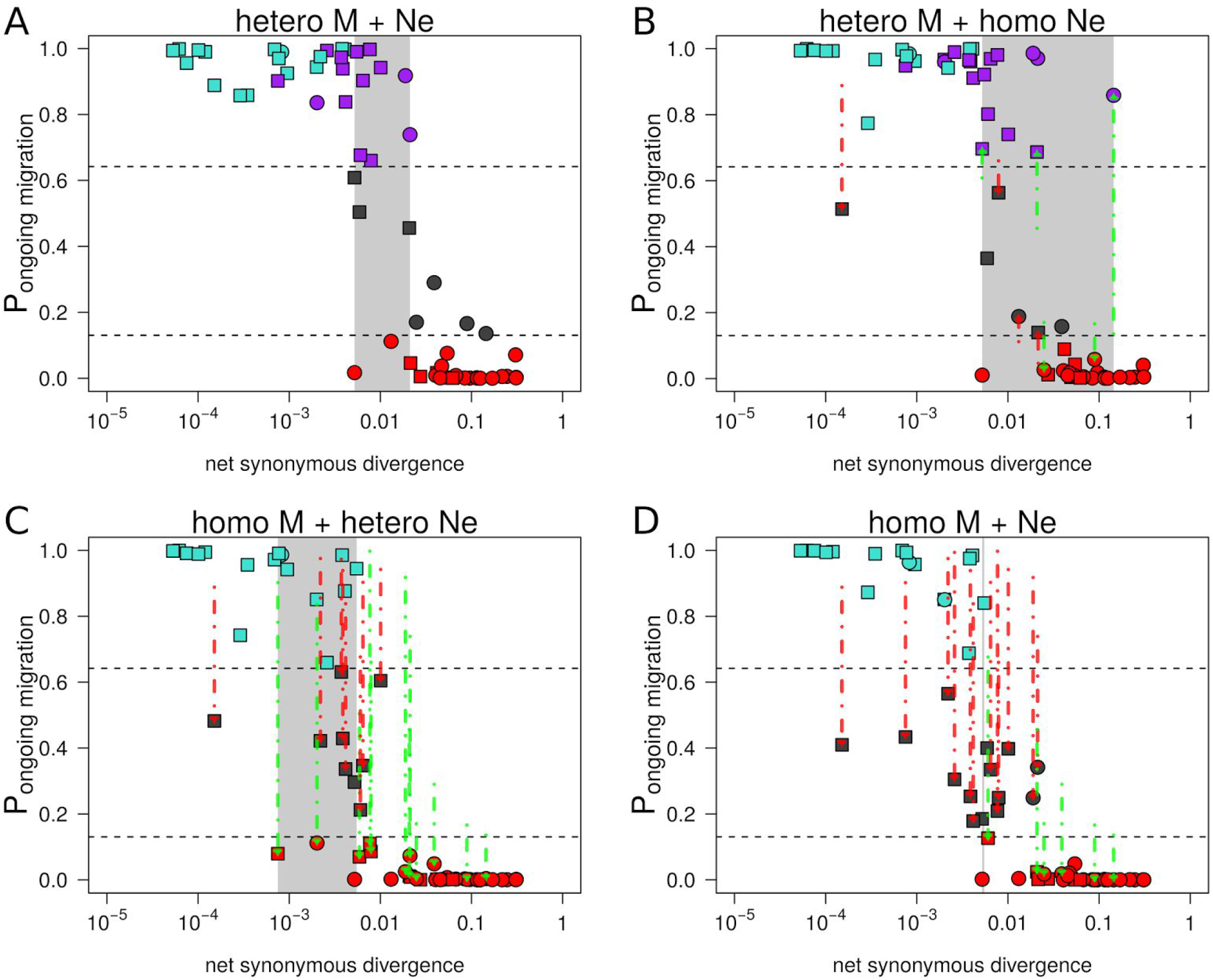
Relation between net synonymous divergence *Da* and probability of ongoing gene flow. Net synonymous divergence is the average proportion of differences at synonymous positions between individuals sampled in the two compared species due to mutations occurring after the ancestral split. The “hetero M + *Ne*” analysis was made by assuming genomic variation for both *M* and *Ne*. The “hetero M” analysis solely takes into account genomic variation in introgression rates over the whole genome. The “hetero *Ne*” analysis solely takes into account genomic variation in *Ne*. The “homo M + *Ne*” analysis considers one value of *M* and one value of *Ne* shared by the whole genome. Red arrows indicate pairs of species inferred as ambiguous in heteroM (robustness < 0.95), heteroNe and homoM_homoN analysis but not in heteroM_heteroN (robustness ≥ 0.95). Green arrows indicate pairs of species with different and unambiguous inferences (robustness ≥ 0.95) made in heteroM, heteroNe and homoM_homoN when compared to heteroM_heteroN.

**Figure S5. Relation between gross synonymous divergence *Dxy* and probability of ongoing gene flow**

Gross synonymous divergence is the average proportion of differences at synonymous positions between individuals sampled in the two compared species, including differences present in the ancestral species.

The “hetero M + *Ne*” analysis was made by assuming genomic variation for both *M* and *Ne*.

The “hetero M” analysis solely takes into account genomic variation in introgression rates over the whole genome.

The “hetero *Ne*” analysis solely takes into account genomic variation in *Ne*.

The “homo M + *Ne*” analysis considers one value of *M* and one value of *Ne* shared by the whole genome.

Red arrows indicate pairs of species inferred as ambiguous in heteroM (robustness < 0.95), heteroNe and homoM_homoN analysis but not in heteroM_heteroN (robustness ≥ 0.95).

Green arrows indicate pairs of species with different and unambiguous inferences (robustness ≥ 0.95) made in heteroM, heteroNe and homoM_homoN when compared to heteroM_heteroN.

**Figure S6.**
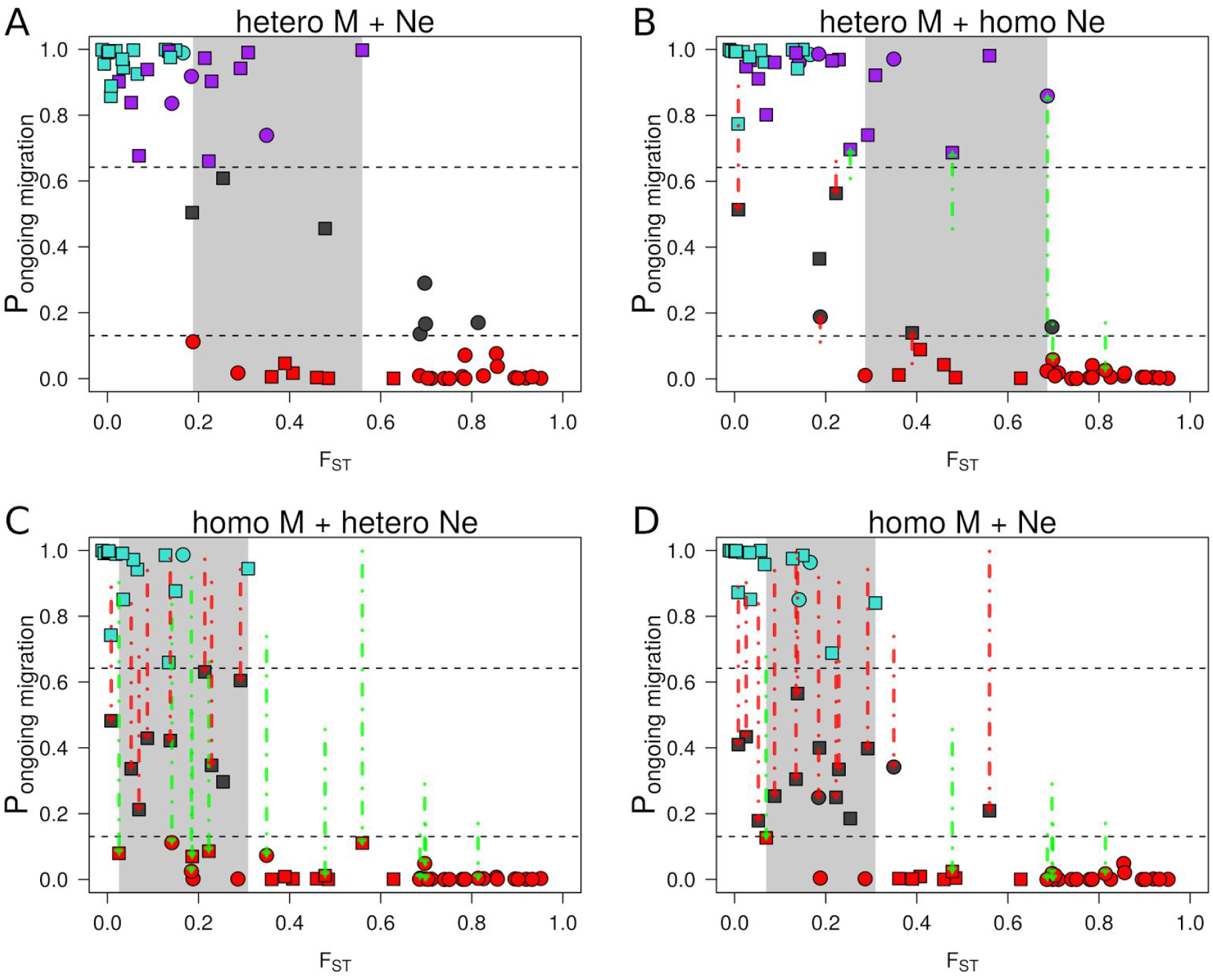
Relation between F_ST_ and probability of ongoing gene flow. The “hetero M + *Ne*” analysis was made by assuming genomic variation for both *M* and *Ne*. The “hetero M” analysis solely takes into account genomic variation in introgression rates over the whole genome. The “hetero *Ne*” analysis solely takes into account genomic variation in *Ne*. The “homo M + *Ne*” analysis considers one value of *M* and one value of *Ne* shared by the whole genome. Red arrows indicate pairs of species inferred as ambiguous in heteroM (robustness < 0.95), heteroNe and homoM_homoN analysis but not in heteroM_heteroN (robustness ≥ 0.95). Green arrows indicate pairs of species with different and unambiguous inferences (robustness ≥ 0.95) made in heteroM, heteroNe and homoM_homoN when compared to heteroM_heteroN.

**Figure S7.**
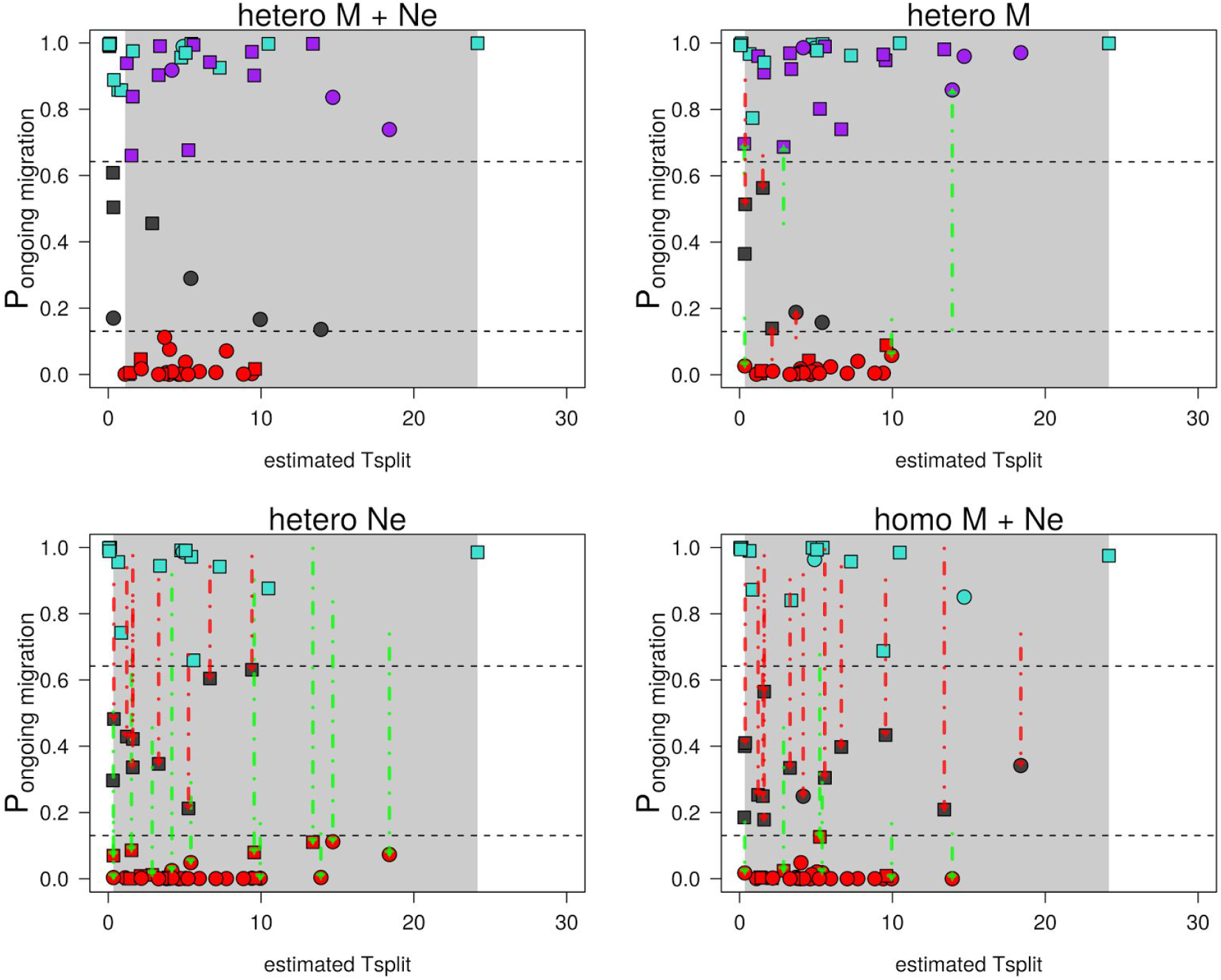
Relation between the estimated *T*_*split*_ under the IM model and probability of ongoing gene flow. The “hetero M + *Ne*” analysis was made by assuming genomic variation for both *M* and *Ne*. The “hetero M” analysis solely takes into account genomic variation in introgression rates over the whole genome. The “hetero *Ne*” analysis solely takes into account genomic variation in *Ne*. The “homo M + *Ne*” analysis considers one value of *M* and one value of *Ne* shared by the whole genome. Red arrows indicate pairs of species inferred as ambiguous in heteroM (robustness < 0.95), heteroNe and homoM_homoN analysis but not in heteroM_heteroN (robustness ≥ 0.95). Green arrows indicate pairs of species with different and unambiguous inferences (robustness ≥ 0.95) made in heteroM, heteroNe and homoM_homoN when compared to heteroM_heteroN.

**Figure S8.**
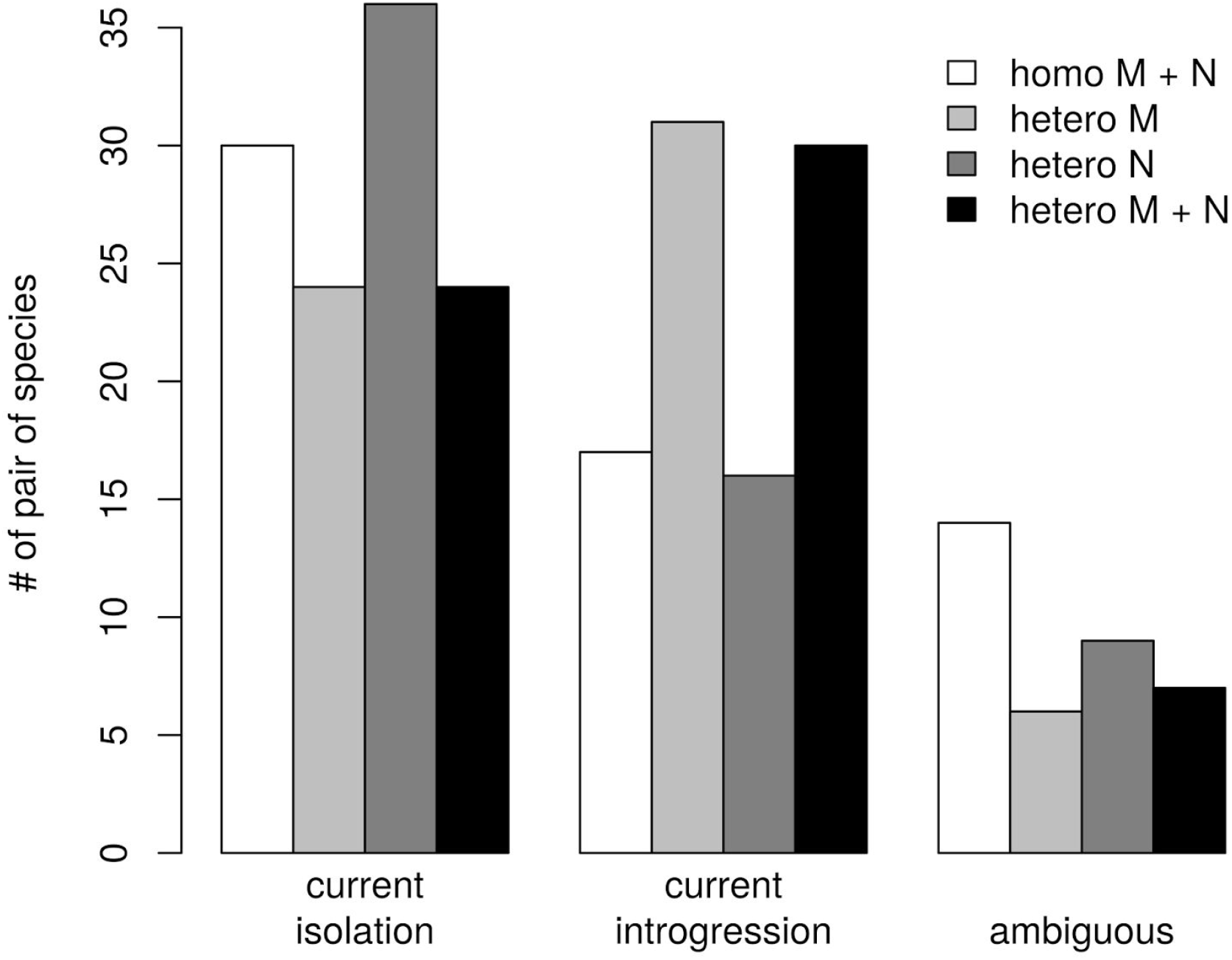
Number of pair of species supporting current isolation, current introgression or ambiguity in model choice. A pair of species is associated to “current isolation” if the sum of posterior probabilities P(SI) + P(AM) is associated to a robustness ≥ 0.95. A pair of species is associated to “current introgression” if the sum of posterior probabilities P(SC) + P(IM) is associated to a robustness ≥ 0.95. The ambiguous status is attributed to a pair of species when “current isolation” and “current introgression” are not strongly supported. The “homo M + N” analysis was made by assuming an unique genomic introgression rate and an unique *Ne* over the whole genome. The “hetero M” analysis takes into account genomic variation in introgression rates over the whole genome. The “hetero N” analysis takes into account genomic variation in *Ne*. The “hetero M + N” analysis takes into account genomic variation in introgression rates and in *Ne*.

**Figure S9. Number of pair of species showing evidences for SI, AM, IM, SC, PAN or ambiguity in model choice for three distinct ABC analysis.**

A pair of species is associated to SI or AM if its relative posterior probability is greater than 0.8696. A pair of species is associated to IM, SC or PAN tf its relative posterior probability is greater than 0.6419.

The “homo M + N” analysis was made by assuming an unique genomic introgression rate and an unique *Ne* over the whole genome.

The “hetero M” analysis takes into account genomic variation in introgression rates over the whole genome.

The “hetero N” analysis takes into account genomic variation in *Ne*.

The “hetero M + N” analysis takes into account genomic variation in introgression rates and in *Ne*.

**Figure S10.**
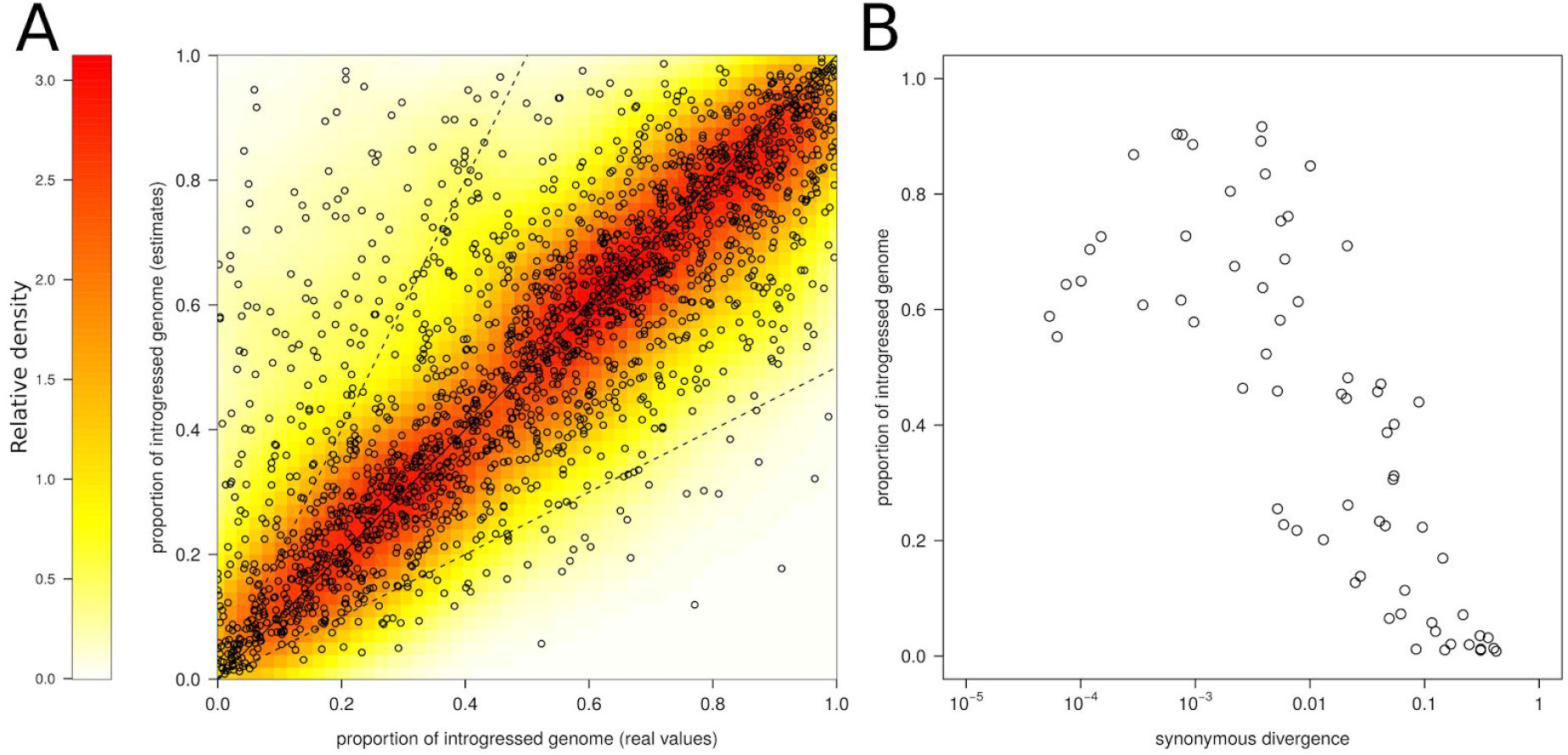
Estimating α, the proportion of loci that introgress, under the IM model. 2,000 pseudo-observed datasets (PODs) were simulated under the IM model with heterogeneity in introgression rates. We estimated the parameters of this model by using the ABC approach described in the ‘Materials and Methods’ section, α is the proportion of the genome crossing the species barrier at a rate *N.m* >0. A. x-axis: values of α used to produce the PODs; y-axis: values of α estimated by ABC from the simulated PODs. Solid line represents f(x) = x. Dotted lines represent f(x) = 2.x and f(x) = x/2 respectively. B. Estimated values of α for the observed pairs of population/species as a function of their net synonymous divergence.

**Figure S11. Estimating *N.m*, the effective migration rate, under the IM model**

2,000 pseudo-observed datasets (PODs) were simulated under the IM model with heterogeneity in introgression rates.

A. x-axis: values of *N.m* used to produce the PODs; y-axis: values of *N.m* estimated by ABC from the simulated PODs.

Solid line represents f(x) = x.

Dotted lines represent f(x) = 2.x and f(x) = x/2 respectively.

B. Estimated values of *N.m* for the observed pairs of population/species as a function of their net synonymous divergence.

**Figure S12.**
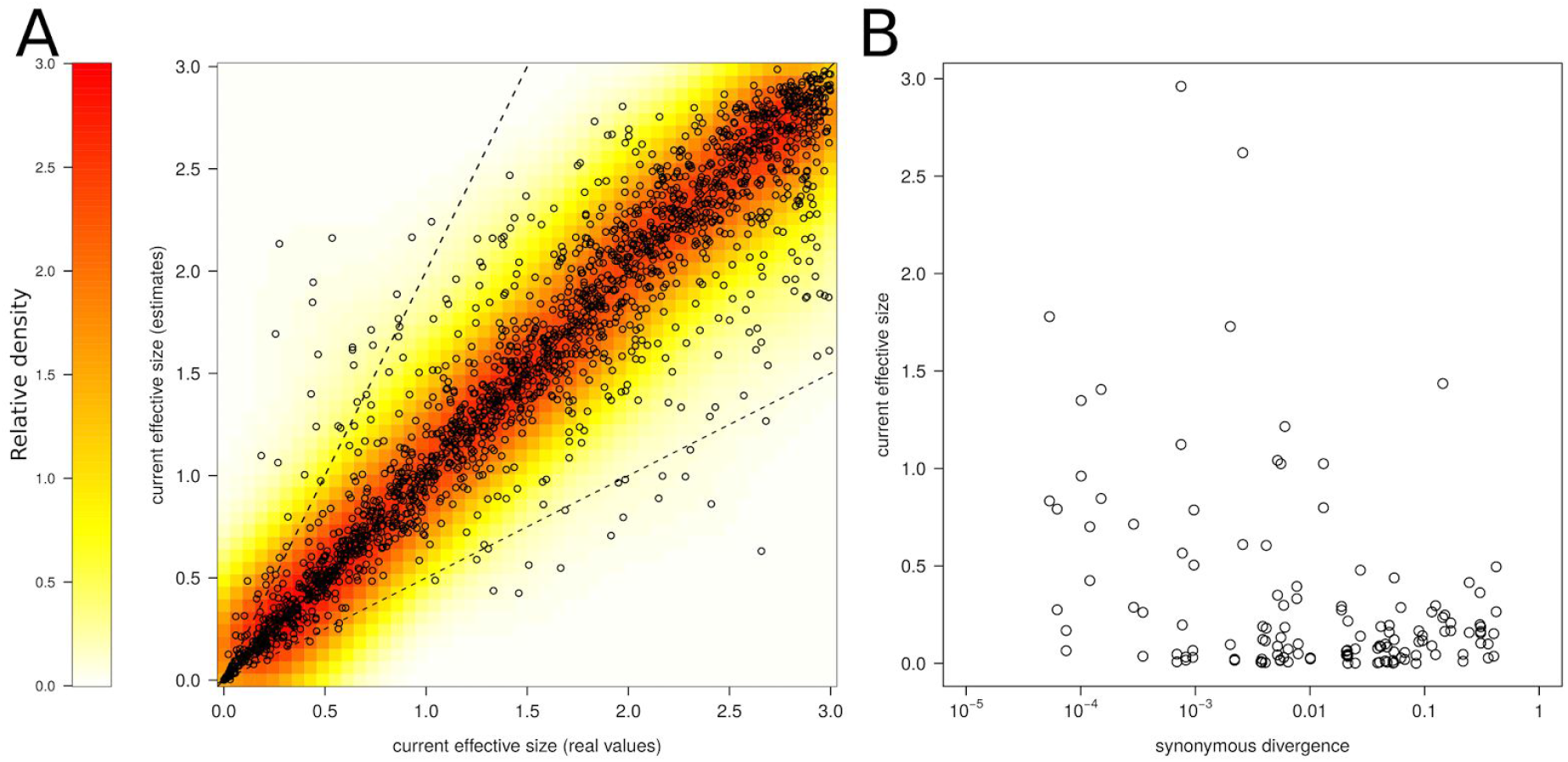
Estimating *N*, the effective population size of daughter populations, under the IM model. 2,000 pseudo-observed datasets (PODs) were simulated under the IM model with heterogeneity in introgression rates. A. x-axis: values of *N* used to produce the PODs; y-axis: current values of *N* estimated by ABC for all PODs. Solid line represents f(x) = x. Dotted lines represent f(x) = 2.x and f(x) = x/2 respectively. B. Estimated values of *N* for the observed pairs of population/species as a function of their net synonymous divergence.

**Figure S13.**
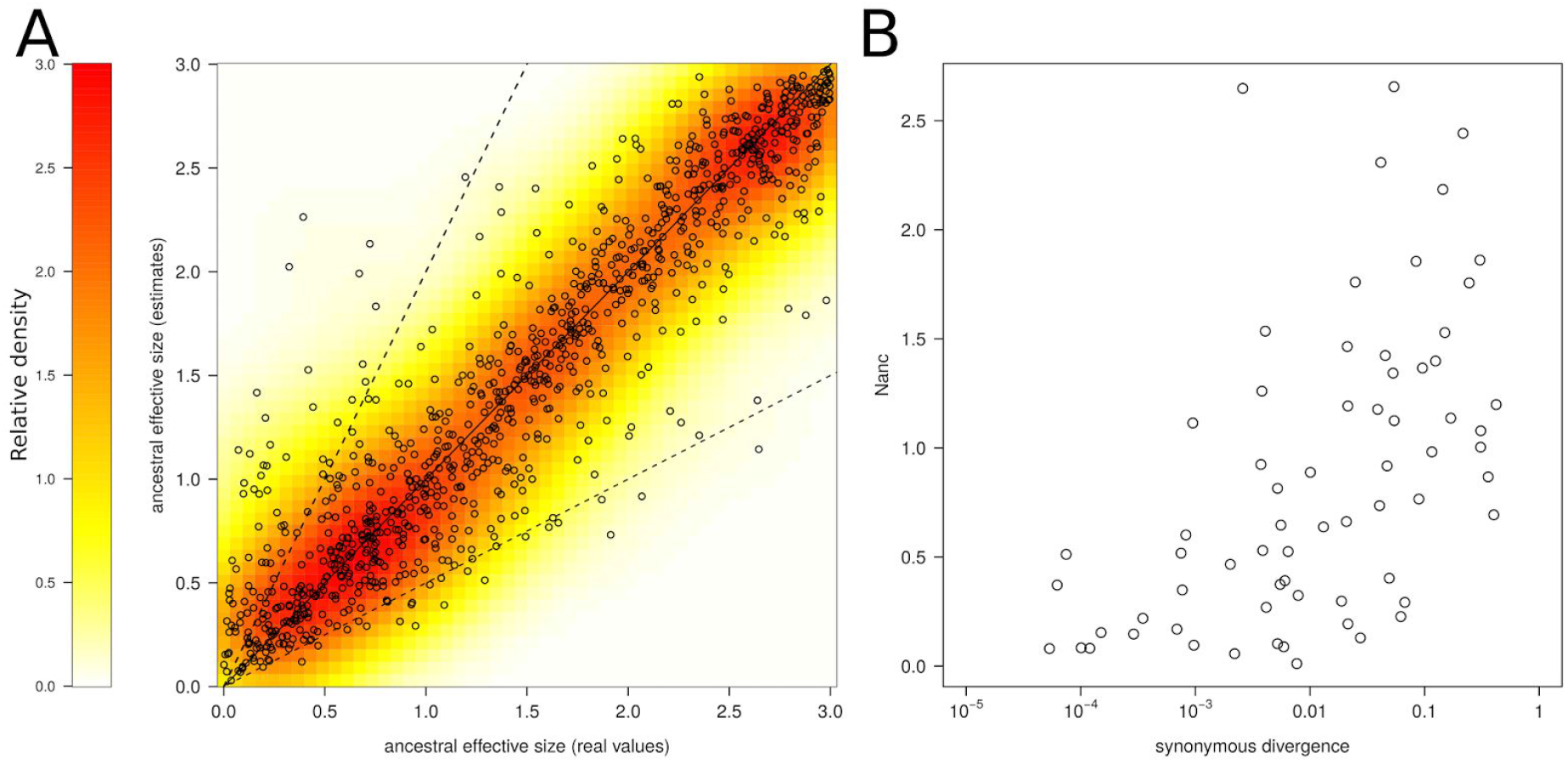
Estimating *Nanc*, the effective size of the ancestral population, under the IM model. 2,000 pseudo-observed datasets (PODs) were simulated under the IM model with heterogeneity in introgression rates. A. x-axis: values of *Nanc* used to produce the PODs; y-axis: estimated values of *Nanc* for all PODs. Solid line represents f(x) = x. Dotted lines represent f(x) = 2.x and f(x) = x/2 respectively. B. Estimated values of *Nanc* for the observed pairs of population/species as a function of their net synonymous divergence.

**Figure S14.**
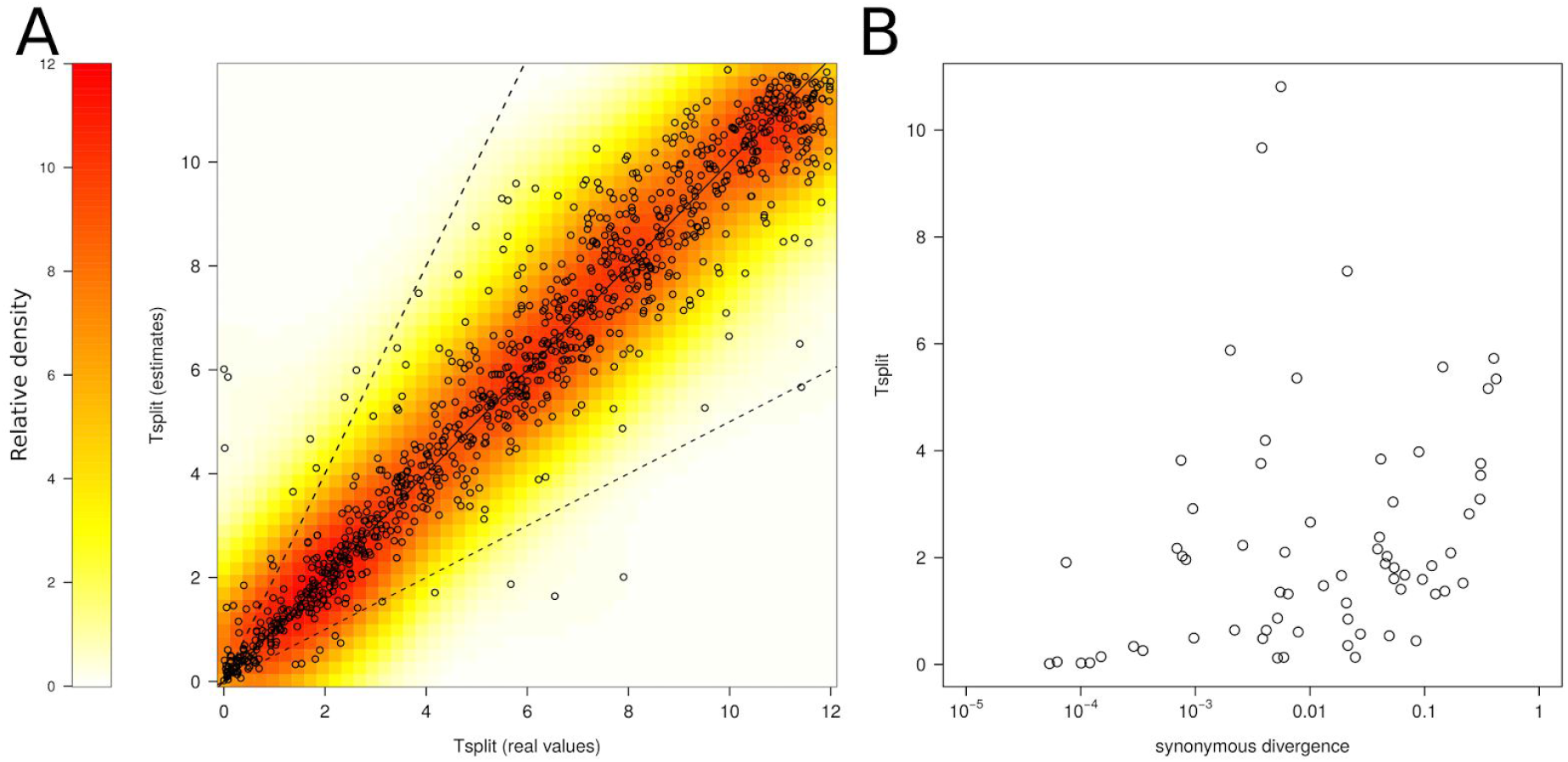
Estimating *T*_*split*_, the time of ancestral subdivision, under the IM model. 2,000 pseudo-observed datasets (PODs) were simulated under the IM model with heterogeneity in introgression rates. *T*_*split*_ is expressed in million of generations since the ancestral separation. A. x-axis: values of *T*_*split*_ used to produce the PODs; y-axis: estimated values of *T*_*split*_ for all PODs. Solid line represents f(x) = x. Dotted lines represent f(x) = 2.x and f(x) = x/2 respectively. B. Estimated values of *T*_*split*_ for the observed pairs of population/species as a function of their net synonymous divergence.

